# Quantifying the efficacy of first aid treatments for burn injuries using mathematical modelling and *in vivo* porcine experiments

**DOI:** 10.1101/131391

**Authors:** Matthew J Simpson, Sean McInerney, Elliot J Carr, Leila Cuttle

## Abstract

First aid treatment of burn injuries reduces scarring and improves healing. Here, we quantify the efficacy of various first aid treatments by using a mathematical model to describe a suite of experimental data from a series of *in vivo* porcine experiments. We study a series of consistent burn injuries that are subject to first aid treatments that vary in both the temperature and duration of the first aid treatment. Calibrating the mathematical model to the experimental data provides estimates of the *in vivo* thermal diffusivity, the rate at which thermal energy is lost to the blood (perfusion), and the heat transfer coefficient controlling the loss of thermal energy at the interface of the fat and muscle layers. A limitation of working with *in vivo* animal experiments is the difficulty of resolving spatial variations in temperature across the tissues. Here, we use the solution of the calibrated mathematical model to predict and visualise the temperature distribution across the thickness of the tissue during the creation of the burn injury and the application of various first aid treatments. Using this information we propose, and report values for, a novel measure of the potential for tissue damage. This measure quantifies two important aspects that are thought to be related to thermal injury: (i) the volume of tissue that rises above the threshold temperature associated with the accumulation of tissue damage; and, (ii) the duration of time that the tissue remains above this threshold temperature. We conclude by discussing the clinical relevance of our findings.

## 1 Introduction

Burn injuries worsen after the initial insult [23], as thermal energy and associated tissue destruction spreads into surrounding tissues. However, if first aid treatment is administered immediately at the scene or prior to qualified medical treatment, burn patients have improved healing outcomes [28]. Optimal first aid treatment conditions have been previously determined using standardised, controlled porcine burn models [4, 9, 10, 29]. Porcine models are used because pig skin is anatomically and physiologically similar to human skin [15, 18], and pig skin responds to therapeutic agents in a similar way to human skin [25]. Recommended first aid treatment is the immediate administration of cool, running water for 20 minutes. This decreases tissue injury, increases the rate of wound healing and reduces scarring [4, 8, 10, 29]. Ethically and morally, there is a limit to the number of different burn and treatment conditions that can be tested using live animals. Therefore, animal experiments cannot be used to examine all potential treatment temperatures and durations.

In this work we use a porcine burn model to study a series of standardised burns created by exposing the surface of the skin to a temperature of 92 °C for 15 seconds. The propagation of thermal energy through the skin is observed by measuring the temperature under the skin using a subdermal temperature probe. Using this approach we examine the effects of various first aid treatments by applying cooling water of various temperatures (0, 2 and 15 °C), for different durations (10, 20, 30 and 60 minutes). This data set provides a wealth of information about how thermal energy propagates through the skin, and clearly illustrates that different first aid treatments have a measurable effect. However, taken in isolation, even a high-quality reproducible experimental data set such as this does not provide sufficient information to quantify the efficacy of different first aid treatments as this requires more detailed spatial and temporal information about the propagation of thermal energy through the skin.

To address this gap in our knowledge we examine a suite of experimental data describing the effect of various first aid treatment strategies using a mathematical model that describes the spatial and temporal distribution of thermal energy within the skin [11, 12, 19, 20, 21]. Owing to the geometry of the experiments, we use a one-dimensional model to describe the temperature of the skin as a function of depth, *x*, and time *t*. The model incorporates three key mechanisms: (i) conduction of thermal energy through the skin; (ii) potential loss of thermal energy to the blood (perfusion); and, (iii) potential loss of thermal energy at the lower boundary, into the deeper muscle tissues. These three processes are characterised by three constant parameters. Calibrating the solution of the model to the experimental data provides estimates of these three parameters. Once calibrated, we are able to use the model to examine the details of the spatial and temporal variation in temperature within the skin to reveal additional details that are not discernible in the experimental design. Using these details, we propose a novel measure to quantify the potential for tissue damage during the experiment. The new measure of potential damage explicitly accounts for the volume of tissue that rises above a threshold temperature, and the duration of time that the tissue remains above that threshold temperature. We calculate this measure of potential damage for a range of first aid treatments to reveal, for the first time, quantitative insight into the efficacy of different first aid treatment options.

## 2 Experimental methods

### 2.1 Animal experimental data

Animal studies were ethically approved (UQAECP&CH/202/06, UQAECMED/RCH/376/08) and all animals were treated humanely and in accordance with the Australian code of practice for the care and use of animals for scientific purposes. The data presented was initially obtained as part of other studies [8, 10]. Data from a total of 37 Large White juvenile pigs are included. The animals were approximately 15-20 kg, or 8 weeks old, when data was collected.

Animals were under a general anaesthetic for all procedures and received appropriate analgesia to minimise any suffering. Anaesthesia was inducted with an intramuscular dose of 13 mg/kg ketamine hydrochloride (Ketamine 100 mg/mL, Parnell Laboratories, Alexandria, Australia) and 1 mg/kg xylazine (Xylazil 20 mg/mL, Ilium, Troy Laboratories, Sydney, Australia) and maintained with halothane via a size 4 laryngeal mask airway. Hair was clipped from the back and flanks, and the skin rinsed with clean water before wounding. Buprenorphine hydrochloride at 0.01 mg/kg (Temgesic 0.3 mg/mL, Reckitt Benckiser, West Ryde, Australia) was administered as an analgesic on induction. The subdermal temperature was measured during burn creation and first aid treatment via a temperature probe inserted under the skin. A 14G 2.1 × 45 mm cannula was obliquely inserted from outside the wound area and advanced through the fat layer under the dermis until the tip was in the centre of the burn area. The needle was removed from the cannula and a type K thermocouple (Radiospares Components Pty Ltd., Smithfield, Australia) was inserted and taped into position. This insertion method was used to minimise the possibility of direct heat transfer onto the probe from the burning device and to minimise damage to the tissue. A digital 54II Fluke thermometer (Fluke Australia Pty Ltd., North Melbourne, Australia) logged temperature measurements every 15 seconds once the burning device was applied and during the course of the first aid treatment. One wound was created in the middle of each flank using a technique described previously [7]. Briefly, a Pyrex laboratory Schott (Mainz, Germany) Duran 500 mL, 75 mm diameter bottle was used (Figure 1a) which had the glass bottom removed and replaced with plastic wrap secured with tape. A volume of 300 mL sterile water was microwaved until it was approximately 95 °C. The temperature of the water inside the bottle was monitored with a digital thermometer (N19-Q1436 Dick Smith, Australia, range 50 to 100 °C). When the water was at 92 °C, the device was placed on the mid flank for 15 seconds.

**Figure 1:**
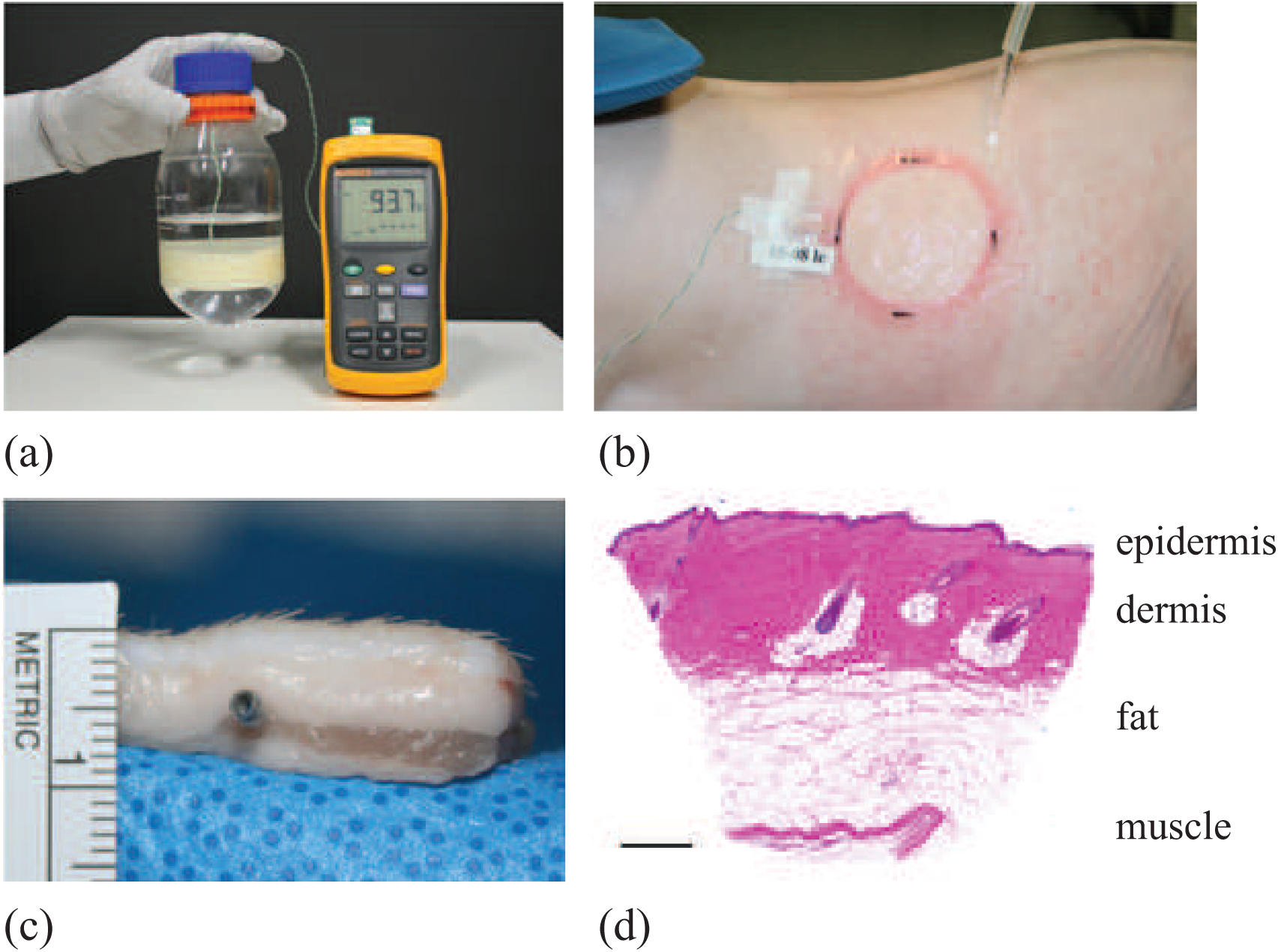
(a) Scalding device. (b) Application of cool running water to the burn with subdermal temperature probe recording the temperature measurements. (c) Excised skin showing the location of the subdermal probe, approximately 4.0 mm below the surface of the skin, and 2.6 mm below the white dermis. (d) Histological image of normal porcine skin. Scale bar corresponds to 1 mm.

### 2.2 Administration of first aid treatment

Treatment commenced as soon as possible after the burn was created. The following different temperature treatments were applied to 25 animals for 20 minutes:

- cool running water at 15 °C;
- cool running water at 2 °C; and,
- granulated ice at 0 °C.

Cool running water was applied at a rate of 1.6 L/min to the wound (Figure 1b) from a temperature controlled waterbath (Grant S26, Jencons Scientific Limited, Leighton Buzzard, UK) to maintain the treatment at a constant temperature. In addition to varying the temperature of the first aid treatments, we also applied 15 °C cool running water to 20 animals for various durations, including:

- 10 minutes;
- 20 minutes;
- 30 minutes; and,
- 60 minutes.

During extended treatment periods, the anaesthetised animals were warmed with hot water bottles and core temperatures were monitored.

### 2.3 Measurement of probe depth

To determine the depth of the temperature probe within the skin, ultrasound imaging (LOGIQe with a L10-22MHz-RS Ultra High Frequency Transducer, GE Healthcare) was used on a different set of animals of similar weight and age with the temperature probe inserted using the same technique. The probes were found to be sitting in the fat layer, 2.6 ± 0.6 mm below the dermis (*n* = 29 wounds). Additionally, a section of tissue with a probe *in situ* was dissected out to macroscopically visualise the location of the probe placement (Figure 1c). A biopsy of normal skin which was fixed in formalin, paraffin embedded, sectioned and stained with Haemotoxylin and Eosin was also obtained as a comparison of the macroscopic and histological appearance of the skin (Figure 1d). The total thickness of the skin (epidermis and dermis) was determined through histological assessment of biopsies collected on the day of treatment from similar weight animals [1], and values adjusted for the distortion created by excision and histological processing to relate it to *in vivo* skin thickness where the skin is stretched [2]. The total skin thickness of the epidermis and dermis was estimated to be 1.4 mm. In summary, the depth of the probe under the surface of the skin is approximately 4.0 mm. Naturally, this depth would vary slightly from animal to animal, and between different locations on the same animal. We will deal with this variability by treating the depth of the skin as a constant value of 4.0 mm, but then averaging our experimental measurements over data from many identically prepared experimental burns.

## 3 Modelling methods

To model the transfer of heat through skin we use a modified one-dimensional Pennes bioheat equation [19]. This model is a linear heat equation with a source term to account for heat loss to the blood supply (perfusion) [11]. The temporal and spatial distribution of temperature, *T* (*x, t*), is governed by,

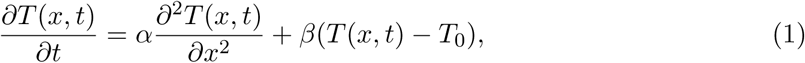

for 0 < *x* < *H* and *t* > 0, subject to the following initial and boundary conditions:

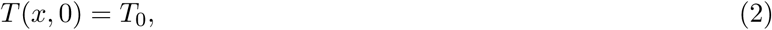

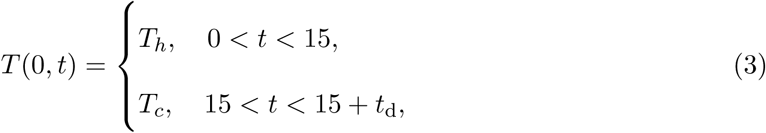

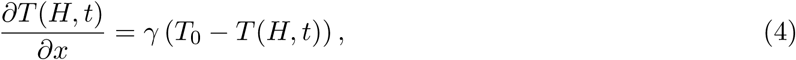

where *H* is the total depth of the tissue including the epidermis, dermis and fat, *t*_*d*_ is the duration for which the treatment is applied, *α* is the thermal diffusivity, *β* is a heat transfer coefficient governing the rate at which thermal energy is lost to the blood supply, *T*_0_ is a reference temperature which we take to be approximately equal to the temperature of the blood and muscle, *T*_*h*_ is the temperature of the heat source used to create the burn, *T*_*c*_ is the temperature of the first aid treatment, and *γ* is a heat transfer coefficient governing the rate at which thermal energy is lost to the adjacent muscle at *x* = *H*. When we apply the model to our experimental data we adopt units of millimetres, seconds and degrees Celsius for the length, time and temperature scales, respectively.

Our modelling work involves several key assumptions. Since the diameter of the burning device (approximately 75 mm, see Figure 1a-b) is much greater than the depth of the tissue (approximately 4 mm, see Figure 1c-d), it is reasonable to work with a one-dimensional model as the measurements are made under the skin, at approximately the centre of the circular burning device. Therefore, locally the heat transfer is effectively one-dimensional [1]. Another key assumption is that we treat *α* and *β* as constants. Although skin is known to be layered [14] (Figure 1d), our measurements of subdermal temperature, *𝒯* (*x, t*), are obtained at one position only, *x* = *H*. This experimental constraint leads us to treat the layered skin as a homogeneous material with spatially averaged thermal properties (Figure 2a) [6]. If we were to invoke a heterogeneous model of multilayered diffusion [5] we would hope to have access to more experimental data to resolve the temperature profile as a function of position, *𝒯* (*x, t*). However, in practice it is impossible to insert multiple temperature probes into a 4 mm layer of skin without compromising the integrity of the tissue (Figure 1c). Therefore, given this constraint, adopting a homogeneous model is reasonable. To solve Equations (1)–(4) we discretise the partial differential equation in space using a central difference approximation with constant spacing, *δx*. The resulting system of coupled ordinary differential equations is integrated through time using a forward Euler approximation, with constant time steps of duration *δt*. We always choose *δx* and *δt* to give grid-independent solutions.

**Figure 2:**
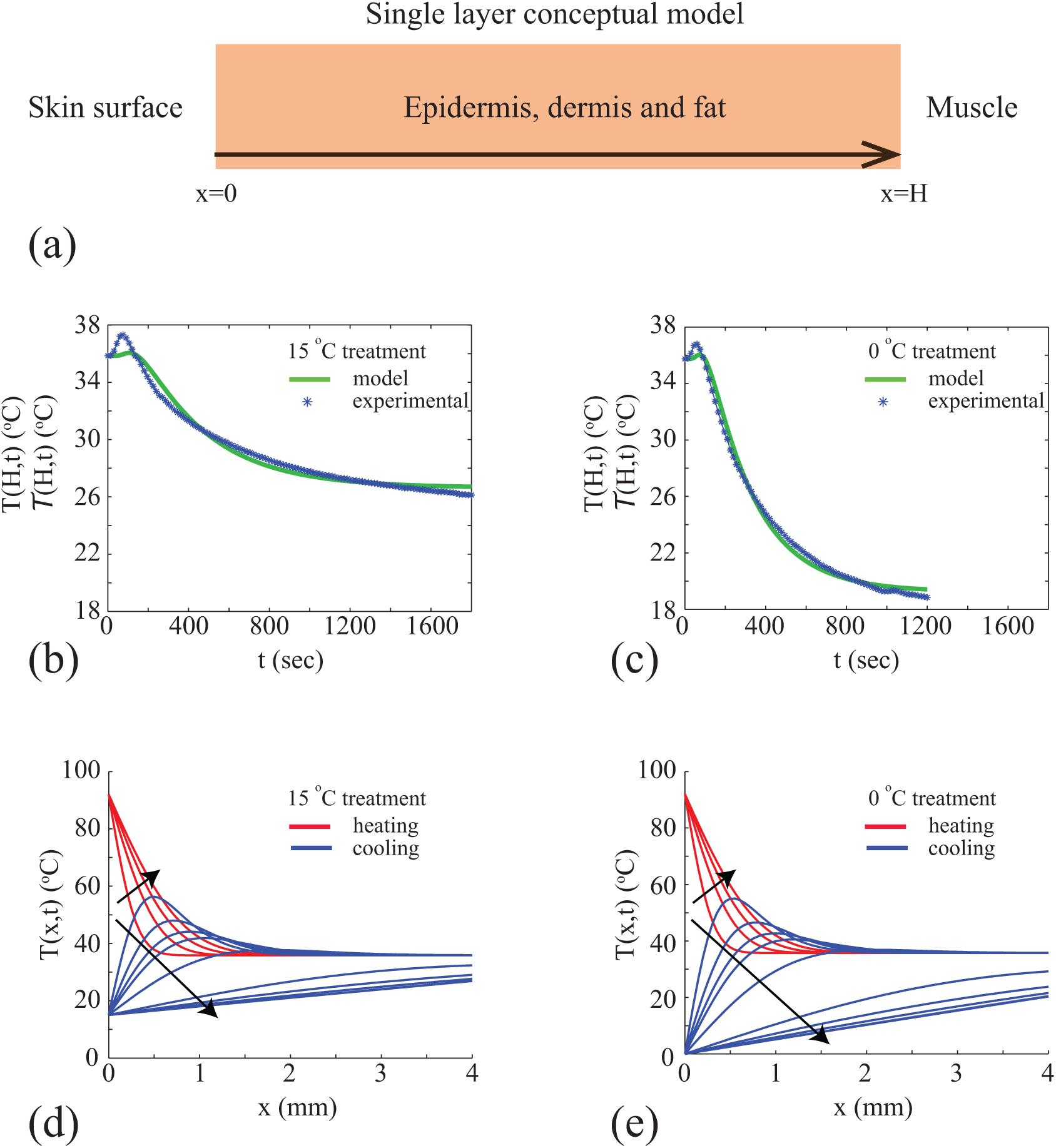
(a) Single homogeneous layer conceptual model. (b)-(c) Calibration of mathematical model to average temperature data for two different first aid treatments with parameters: (*α* = 0.01, *β* = 0.00 and *γ* = 0.32), and (*α* = 0.02, *β* = 0.00 and *γ* = 0.29), respectively. Numerical solutions are obtained with *δx* = 0.1 mm and *δt* = 0.1 seconds. (d)-(e) Solution of calibrated mathematical model showing spatial and temporal variations in the temperature distribution during both the heating (red) and cooling (blue) phases of experiments for two different treatment conditions. Numerical solutions are obtained with *δx* = 0.05 mm and *δt* = 0.0125 seconds. Profiles in (d) are shown at 4 second intervals from *t* = 2 until *t* = 30 seconds, and then at 230 second intervals from *t* = 50 until *t* = 1200 seconds. Profiles in (e) are shown at 4 second intervals from *t* = 2 until *t* = 30 seconds, and then at 250 second intervals from *t* = 50 until *t* = 1800 seconds. In both (d) and (e) the direction of increasing *t* is shown with the arrows. Profiles in (d) and (e) corresponding to the heating phase, 0 < *t* < 15 seconds, are shown in red. Profiles in (d) and (e) corresponding to the cooling phase, *t* > 15 seconds, are shown in blue.

In this study we treat certain parameters in the mathematical model as known, either through making explicit experimental measurements, or invoking reasonable assumptions. Known parameters include:

1. the depth of tissue, *H* = 4.0 mm;
2. the initial temperature of the tissue, *T*_0_ = 36 °C;
3. the temperature of the heat source used to create the burn, *T*_*h*_ = 92 °C;
4. the temperature of the first aid treatment, *T*_*c*_ = 0, 2 or 15 °C; and,
5. the duration of the first aid treatment, *t*_d_ = 600, 1200, 1800 or 3600 seconds.

Measurements of the subdermal temperatures, *𝒯* (*H, t*), are recorded during all experiments, and this data is available in the Supplementary material document. To estimate the unknown parameters, the numerical solution of Equations (1)–(4), *T* (*H, t*), is calibrated to experimental measurements, *𝒯* (*H, t*). The calibration procedure involves finding values of *α*, *β* and *γ* so that the solution of Equations (1)–(4) at *x* = *H*, namely *T* (*H, t*), matches the measured data, *𝒯* (*H, t*). The calibration is performed using MATLAB’s nonlinear least–squares routine [13]. In each case we always take care to check that the least–squares parameter estimates are insensitive to the initial choice required for the iterative least–squares algorithm to converge [13].

As a way of quantitatively assessing the effectiveness of the different treatments, we propose the following expression to quantify the potential damage:

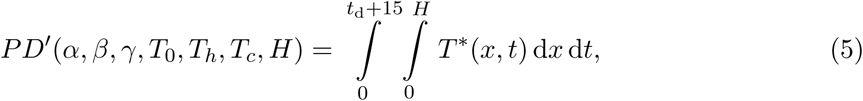

where:

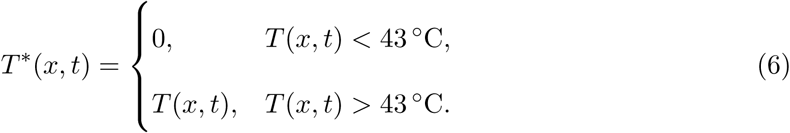

This measure of potential damage, *P D′*, provides a means of quantifying the volume of tissue that experiences temperatures exceeding 43 °C over time. This reflects the fact that tissue damage is thought to occur when tissues are raised above a critical threshold, which is reported to be approximately 43 °C [16, 17]. We remark that the upper limit of integration in Equation (5) for the time variable is written as *t*_d_ + 15, corresponding to the during of the entire experiment, that is the duration of the burn plus the duration of the treatment. However, as we will see later, this upper limit can be truncated well before this time because the integrand, *T* ^*\**^(*x, t*), vanishes well before *t*_d_ + 15.

In reality, *P D ′* depends on all the various parameters in the model. Therefore, strictly speaking, we write *P D ′* (*α, β, γ, T*_0_*, T*_*h*_*, T*_*c*_*, H*) in Equation (5). However, in any practical application of the model we have no control over some of these parameters. For example, values of *α, β, γ* and *T*_0_ cannot to be varied by changing the design of the first aid treatment. Two key variable of interests in this study are the temperature of the cooling mechanism and the duration of the first aid treatment. Therefore, from this point we write *P D ′* (*T*_*c*_*, t*_d_). To further simplify the presentation of our results we use a non-dimensional format,

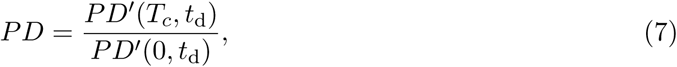

meaning that *P D* is the potential damage function relative to the potential damage incurred by treating the burn with granulated ice, at 0 °C applied for a duration of *t*_d_ seconds.

## 4 Results and discussion

We consider six different first aid treatments, and we refer to these as six different experimental designs because each first aid treatment involves altering a particular variable, such as *t*_d_ and *T*_*c*_. The features of the six experimental designs are summarised in Table 1. For each experimental design, different numbers of experimental replicates were performed. The number of experimental replicates, *n*, for each design is given in Table 1. In brief, each experimental design is assessed using at least *n* = 8 independent, identically created experiments.

**Table 1:** Experimental design. Data showing the number of experimental replicates, *n*, for each experimental design.

| $T_c$ | $t_d = 600$ seconds | $t_d = 1200$ seconds | $t_d = 1800$ seconds | $t_d = 3600$ seconds |
| --- | --- | --- | --- | --- |
| 0 °C | 0 | 18 | 0 | 0 |
| 2 °C | 0 | 16 | 0 | 0 |
| 15 °C | 8 | 16 | 8 | 8 |

The experimental data, *𝒯* _*k*_(*H, t*), for *k* = 1, 2, 3*,… n*, are given in the Supplementary material document for each experimental design. To minimise the effects of experimental variability, we calculate the average experimental data for each experimental design,

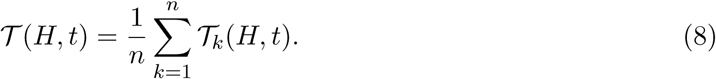

Averaged experimental data, *𝒯* (*H, t*), are also given in the Supplementary material document. When calibrating the solution of Equations (1)-(4) to match the experimental data, we use the averaged data, *𝒯* (*H, t*), to account for variability between the different experimental replicates. For each experimental design we obtain least–squares estimates of *α*, *β* and *γ*, as summarised in Table 2. Since we have six experimental designs, the calibration procedure gives six values of each parameter, and we report the sample mean and sample standard deviation in Table 2. Individual estimates of *α*, *β* and *γ* for each experimental design are reported in the Supplementary material document. Results in Table 2 reveal several insightful trends. For example, the estimate of the thermal diffusivity, *α* = 0.014 mm^2^/s, is consistent with previously determined estimates in a separate study [1], and the variability in the estimates of *α* among the six different experimental designs is very low. For example, the sample standard deviation for *α* is an order of magnitude smaller than the sample mean, giving a coefficient of variation of approximately 28%. This degree of variability is extremely small compared to the reported variability in estimates of diffusivities in biological contexts [24, 26]. Another insightful result is that the estimate of *β* is extremely small. This means that our model calibration procedure suggests that perfusion of thermal energy into the blood supply is negligible in these experiments. Furthermore, our estimates of *β* and *γ* suggest that loss of thermal energy from the system is dominated by heat transfer to the muscle layer at *x* = *H* rather than to perfusion into the blood.

**Table 2:** Parameter estimates, showing the sample mean and sample standard deviation for *α*, *β* and *γ* using the six estimates of each parameter associated with the six different experimental designs.

| Parameter | Units | Sample mean | Sample standard deviation |
| --- | --- | --- | --- |
| $\alpha$ | $\text{mm}^2/\text{s}$ | 0.014 | 0.004 |
| $\beta$ | $1/\text{s}$ | $9.1 \times 10^{-8}$ | $9.2 \times 10^{-8}$ |
| $\gamma$ | $1/\text{mm}$ | 0.32 | 0.08 |

Given our estimates of *α*, *β* and *γ* for each experimental design (Supplementary material), we compare *T* (*H, t*) and *𝒯* (*H, t*) for two different first aid treatment conditions in Figure 2b-c. Results in Figure 2b, for *T*_*c*_ = 15 ° C and *t*_d_ = 1800 seconds, show that the calibrated solution of the model captures the main feature of the *𝒯* (*H, t*) data. This includes both the initial relatively rapid rise in temperature as a result of the application of the burn, as well as the slower cooling over a longer period of time when the cooling treatment is applied at the surface. Results in Figure 2c, for *T*_*c*_ = 0 ° C and *t*_d_ = 1200 seconds, also shows that the calibrated solution of the model captures the main features of the measured *𝒯* (*H, t*) data. Comparing results in Figures 2b-c we clearly see the impact of the different first aid treatments as *T* (*H, t*) reduces to a lower temperature, and at a faster rate when the 0 ° C treatment is applied compared to the 15 ° C treatment. Since the least–squares estimates of *α*, *β* and *γ* do not vary too much between the different experimental designs (Table 1), from this point forward we will use the average values of these parameters as reported in terms of the sample mean in Table 2.

An interesting feature of our experimental data set is that the surface temperature rises to 92 ° C at the surface of the skin, at *x* = 0, during the first 15 seconds of the experiment. However, the temperature at the base of the tissue, at *x* = *H*, never rises above approximately 38 ° C. This highlights one of the limitations of taking a purely experimental approach. As tissue damage is thought to occur when the temperature rises above 43 ° C [16, 17], one of the key aspects of interest is to examine the complete spatial and temporal distribution of temperature throughout the tissue. Understanding how the temperature varies across the skin layers would allow us to quantitatively assess how different first aid treatments affect the temperature of the tissues relative to the damage threshold. Since our experimental data provides information at one depth only, *x* = *H*, it is unclear what the temperature distribution across the entire layer of skin is, and it is unclear how different treatments affect the distribution of temperatures within the skin layer based on the experimental data alone.

Results in Figure 2d-e show *T* (*x, t*) to provide additional insight into the details of the spatial and temporal variation in temperature across the depth of the skin. These results correspond to the experimental designs associated with the results in Figure 2b-c, respectively. The temperature profiles during the first part of the experiment when the burn is created, 0 < *t* < 15 seconds, is identical in both cases. However, the temperature profiles differ during the latter part of the experiment, *t* > 15 seconds, where the different first aid treatments are applied. A qualitative comparison of the *T* (*x, t*) profiles in Figure 2d-e indicates that the main differences are in the upper layer of the tissue, close to the surface at *x* = 0 mm, whereas the measured differences are at the base of the tissue, at *x* = 4 mm. Furthermore, the differences between the 15 ° C and 0 ° C treatments becomes more pronounced at later times. This observation suggests that an improved understanding of the efficacy of the various first aid treatments can be obtained by focusing on the details of the temperature distribution across the entire layer of skin, rather than at just one location.

To provide more detail about how the various first aid treatments affect the spatial and temporal variation of temperature within the tissue we plot *T* (*x, t*) using the average values of *α*, *β* and *γ* (Table 2) in a series of space-time diagrams [27] in Figure 3a-c. These plots are constructed for a range of first aid treatment temperatures, *T*_*c*_, including some of the conditions explored experimentally (*T*_*c*_ = 0 and *T*_*c*_ = 15 ° C), as well as other first aid treatment temperatures that extrapolate to conditions that were not examined experimentally (*T*_*c*_ = 25 ° C). All results in Figure 3 correspond to the same duration of treatment, *t*_d_ = 1200 seconds. Exploring the effects of varying *T*_*c*_ can provide very practical insight. For example, results in Figure 3a-b correspond to *T*_*c*_ = 0 and 15 ° C, respectively. These conditions could correspond to treating a burn injury with ice cold water and tap water in a cool climate, respectively. In contrast, results in Figure 3c correspond to *T*_*c*_ = 25 ° C, which could correspond to treating a burn injury with tap water in a warm, tropical environment. In each plot we truncate the *T* (*x, t*) profile, showing only the spatial and temporal distribution for *T* (*x, t*) *≥* 43 ° C as this is thought to be the critical temperature where tissue damage occurs [16, 17].

**Figure 3:**
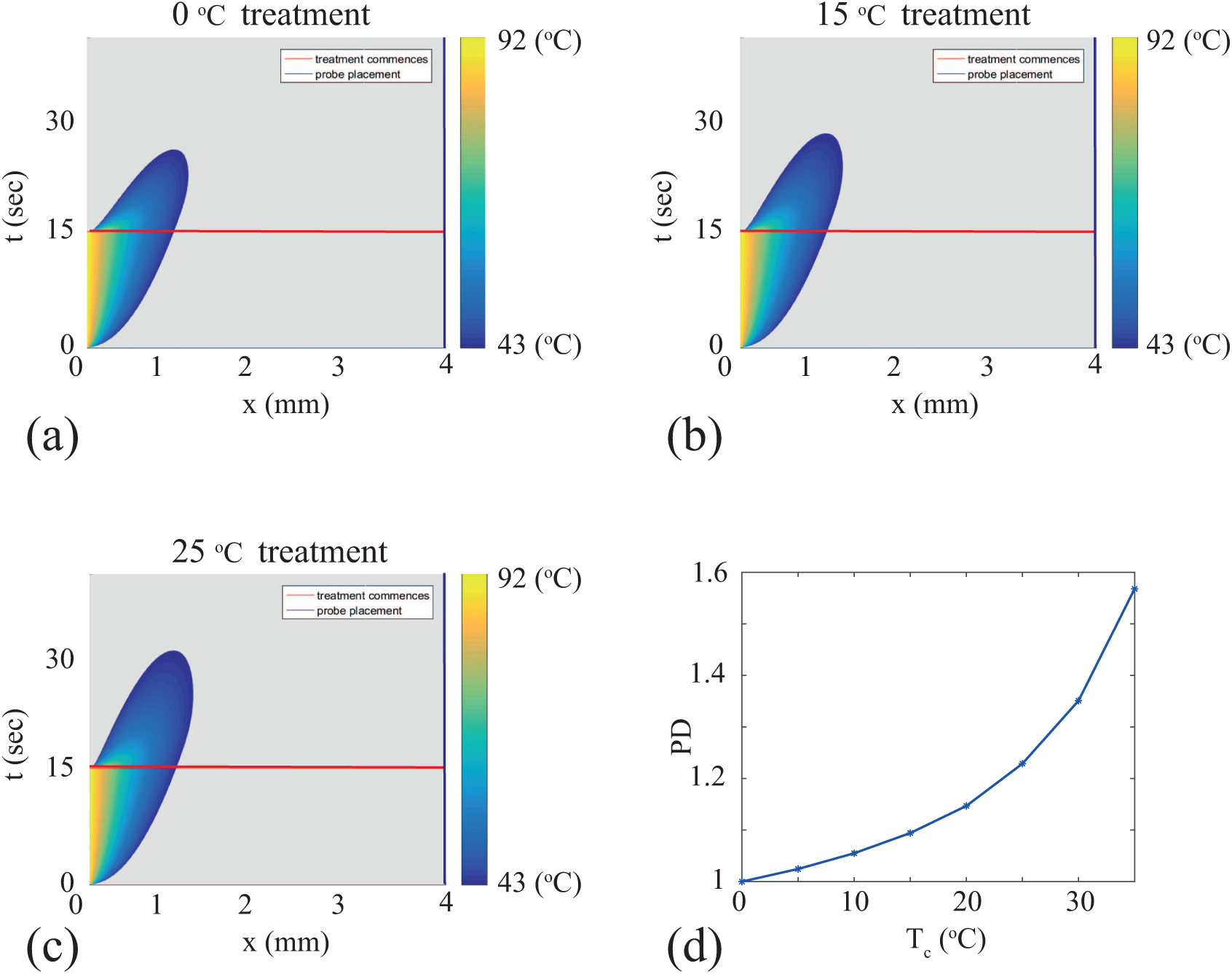
(a)-(c) Space-time diagrams visualising Equation (6), using the numerical solution of Equations (1)–(4) for *T* (*x, t*) with: (a) *T*_*c*_ = 0 ° C; (b) *T*_*c*_ = 15 ° C; and, (c) *T*_*c*_ = 25 ° C. In each case *t*_d_ = 1200 seconds, and all other parameters are given in Table 1. Each space-time diagram shows *T* (*x, t*), for *T* (*x, t*) > 43 ° C as indicated by the color bars, and grey where *T* (*x, t*) < 43 ° C. (d) shows the potential damage, *P D*, given by Equation (7). Values of *P D* are given for *T*_*c*_ increasing from 0 ° C to 35 ° C, in increments of 5 ° C. To calculate *P D* we evaluate the double integral in Equation (5) using the rectangle rule by discretizing the function *T* ^*\**^(*x, t*) with *δx* = 0.05 mm and *δt* = 0.0125 seconds.

Each space-time diagram in Figure 3a-c includes a horizontal line at *t* = 15 seconds. This line denotes the transition from the first part of the experiment where the burn is created, to the latter part of the experiment where the first aid is applied. Since all experiments are initiated with the same burn condition, the space-time diagrams in Figure 3a-c are identical for *t* < 15 seconds. The differences associated with the various first aid treatments is visually discernible in the upper region of the space-time diagrams, where *t* > 15 seconds. Here we see that the spatial and temporal extent of the region where *T* (*x, t*) > 43 ° C reduces as *T*_*c*_ is reduced. Each space-time diagram in Figure 3a-c also includes a vertical line at *x* = 4 mm to denote the depth at which experimental measurements are made in our experimental design. Interestingly, as we noted previously for the results in Figure 2d-e, the temperature at *x* = 4 mm never rises above the threshold of 43 ° C [16, 17]. However, comparing the spatial and temporal extent of the region where *T* (*x, t*) > 43 ° C in Figure 3a-c confirms that different first aid treatment options lead to clear differences in terms of which part of the tissue rises above the threshold of 43 ° C [16, 17], and how long the tissue remains above this threshold. Overall, the main effect is that cooler first aid treatments lead to a reduction in the area of the space-time diagram where *T* (*x, t*) > 43 ° C. While this result is intuitively obvious, the quantitative details of how rapidly this area decreases with decreasing *T*_*c*_ is not obvious from intuition alone.

To provide quantitative insight into how the potential for tissue damage varies with the experimental design, we calculate *P D*, given by Equations (5)-(6), for a suite of first aid treatments, and results are presented in Figure 3d. These results show how *P D*^*\**^ (Equation 7) increases with *T*_*c*_, and the details of this dependence reveals some practical information. For example, since *P D* is an increasing function of *T*_*c*_, there is always a benefit in treating a burn injury with cooler water. However, the slope of the curve in Figure 3d reveals further insight. For example, there is a relatively large reduction in *P D* if the temperature of the treatment water is reduced from 35 ° C to 30 ° C. In this case a 5 ° C reduction in the temperature of the cooling water reduces *P D* from 1.57 to 1.35. However, there is a relatively smaller reduction in *P D* if the temperature of the treatment water is reduced from 5 ° C to 0 ° C. In this case, a 5 ° C reduction in the temperature of the cooling water reduces *P D* from just 1.02 to 1.00. In summary, the benefit of reducing the temperature of the cooling water is most pronounced for modest to warm first aid treatment temperatures.

## 5 Conclusions

In this work we present a combined experimental and mathematical modelling study to quantify the efficacy of various first aid treatments using a suite of data from *in vivo* porcine experiments. In particular, we examine a series of consistent burn injuries that are then subject to various designs of first aid treatments by varying both the temperature and duration of the first aid treatment. Using this extensive experimental data set, we calibrate a mathematical model to reveal estimates of the thermal diffusivity, the rate at which thermal energy is lost to the blood, and the heat transfer coefficient determining the rate at which thermal energy is lost at the lower boundary of the tissue, at the interface of the fat and muscle layers. This calibration exercise provides precise estimates of the *in vivo* thermal diffusivity, and suggests that the loss of thermal energy in the system is dominated by loss to the muscle rather than loss to the blood. Previous mathematical modelling studies have assumed that thermal energy loss to the blood is negligible [11], and here we provide novel *in vivo* evidence to support such assumptions.

The key benefit of working with the mathematical modelling framework is that our extensive experimental data set shows the temperature at just one position at the bottom of the skin tissues, at *x* = 4.0 mm, which is at the interface of the fat and muscle. Although burns are created by exposing the surface of the skin, at *x* = 0 mm, to a temperature of 92 ° C, the experimental data at the base of the skin tissues, *x* = 4.0 mm, does not record any increase in temperature above approximately 38 ° C. This is important because it is thought that damage to tissues occurs when the temperature increases above a threshold temperature of approximately 43 ° C [16, 17] and while we know that the temperature of the surface of the skin rises well above this threshold, all experimental measurements of temperature at the base of the tissue remain below the threshold. Therefore, using the experimental data alone, it is unclear how much of the tissue rises above the threshold temperature, and it is also unclear how long the temperature of the tissue remains above the threshold temperature. In contrast to the experimental data, the solution of the calibrated mathematical model illustrates the temperature distribution right across the thickness of the tissue and shows precisely how different first aid treatments affect the volume of tissue that rises above 43 ° C, and the duration of time that the tissue rises above this threshold. To quantify this we propose a measure of the potential tissue damage, *P D*, and we show how different first aid treatment designs influence this measure.

The focus of this work is to combine an *in vivo* experimental data set and a mathematical model to provide quantitative insight into how first aid treatments affect the spatial and temporal distribution of temperature within living skin. Developing a quantitative understanding of the heat transfer process is an essential first step before we can attempt to develop a quantitative understanding of how first aid treatments affect tissue damage. Therefore, this study focuses on the heat transfer processes only. As a consequence, the modelling presented here indicates that first aid treatment is likely to be more beneficial if the treatment water is cooler. One might be tempted to conclude from this study that applying ice at 0 ° C is the optimal first aid treatment. However, previous investigations focusing on tissue damage and healing outcomes suggest that ice must never be applied to a burn injury as it is not beneficial for healing [9], may lead to additional tissue damage [22], and might also increase the risk of hypothermia. However, our study suggests that if tap water in a tropical climate is warmer than 25 ° C, it may be beneficial to apply cooler refrigerated water rather than local tap water for maximum effect. We suggest that further studies that aim to quantify how first aid treatments affect spatial and temporal tissue damage are warranted, and we leave this as a topic for future investigation.

## Author Contributions

MS jointly coordinated the research project, formulated the mathematical model, supervised the development and application of numerical algorithms, and drafted the manuscript. SM developed and applied the numerical algorithms to the experimental data. EC formulated the mathematical model, supervised the development and application of numerical algorithms, and helped draft the manuscript. LC jointly coordinated the research project, generated the experimental data, and helped draft the manuscript. All authors gave final approval for publication.

## Acknowledgements

This work is supported by the Australian Research Council (DE150101137, DP170100474), the National Health and Medical Research Council (APP1130862, APP409902) and the Australian Mathematical Sciences Institute who provided SM with a 2016-2017 Vacation Research Scholarship.

